# MED19 alters AR occupancy and gene expression in prostate cancer cells, driving MAOA expression and growth under low androgen

**DOI:** 10.1101/857235

**Authors:** Hannah Weber, Rachel Ruoff, Michael J. Garabedian

**Affiliations:** Departments of Microbiology and Urology, NYU School of Medicine, New York, NY, USA

## Abstract

Androgen deprivation therapy (ADT) is a mainstay of prostate cancer treatment, given the dependence of prostate cells on androgen and the androgen receptor (AR). However, tumors become ADT-resistant, and there is a need to understand the mechanism. One possible mechanism is the upregulation of AR co-regulators, although only a handful have been definitively linked to disease. We previously identified the Mediator subunit MED19 as an AR co-regulator, and reported that MED19 depletion inhibits AR transcriptional activity and growth of androgen-insensitive LNCaP-abl cells. Therefore, we proposed that MED19 upregulation would promote AR activity and drive androgen-independent growth. Here, we show that stable overexpression of MED19 in androgen-dependent LNCaP cells promotes growth under conditions of androgen deprivation. To delineate the mechanism, we determined the MED19 and AR transcriptomes and cistromes in control and MED19 LNCaP cells. We also examined H3K27 acetylation genome-wide. MED19 overexpression selectively alters AR occupancy, H3K27 acetylation, and gene expression. Under conditions of androgen deprivation, genes regulated by MED19 and genomic sites occupied by MED19 and AR are enriched for ELK1, a transcription factor that binds the AR N-terminus to promote select AR-target gene expression. Strikingly, MED19 upregulates expression of monoamine oxidase A (MAOA), a factor that promotes prostate cancer growth. MAOA depletion reduces androgen-independent growth. MED19 and AR occupy the MAOA promoter, with MED19 overexpression enhancing AR occupancy and H3K27 acetylation. Furthermore, MED19 overexpression increases ELK1 occupancy at the MAOA promoter, and ELK1 depletion reduces MAOA expression and androgen-independent growth. This suggests that MED19 cooperates with ELK1 to regulate AR occupancy and H3K27 acetylation at MAOA, upregulating its expression and driving androgen independence in prostate cancer cells. This study provides important insight into the mechanisms of prostate cancer cell growth under low androgen, and underscores the importance of the MED19-MAOA axis in this process.

**Author summary:** Prostate cancer is one of the most common cancers worldwide, and androgen hormones are essential for prostate cancer growth. Androgens exert their effects through a protein called the androgen receptor (AR), which turns on and off genes that regulate prostate cancer growth. Powerful drugs that block AR action by lowering androgen levels – so-called androgen deprivation therapy - are used to treat prostate cancer patients, and these yield initial success in reducing tumor growth. However, over time, tumors circumvent androgen deprivation therapy and patients relapse; in many cases, this occurs because AR becomes re-activated. The factors responsible for re-activating AR and promoting growth under androgen deprivation are not well understood. Here, we demonstrate that a subunit of the Mediator transcriptional regulatory complex, called MED19, promotes growth of prostate cancer cells under low androgen conditions, mimicking the ability of tumors to grow under androgen deprivation in prostate cancer patients. MED19 promotes androgen-independent growth by working with a transcription factor that interacts with AR, called ELK1, to induce the expression of genes regulated by AR that promote prostate cancer growth. This study provides important insight into how prostate cancer cells can maintain growth under androgen deprivation through MED19.

## Introduction

Prostate cells depend on androgens and the androgen receptor (AR) for growth and survival, and AR is a key driver of prostate cancer from early to late stage disease [1]. The mainstay of prostate cancer treatment is androgen deprivation therapy (ADT), to which patients initially respond [2–4]. However, AR re-activation occurs following ADT, giving rise to castration-resistant prostate cancer (CRPC) that can grow in the face of low circulating androgens. Although current treatments, such as enzalutamide and abiraterone, extend survival in CRPC patients, none are curative [2, 4–6]. There is a pressing need to better understand the molecular mechanisms behind AR re-activation following ADT.

One important mechanism of AR re-activation with the potential to advance treatment modalities is the upregulation of AR co-regulators [1, 7–9]. For example, CBP (CREB binding protein) and p300, both histone acetyl transferases (HATs), are known AR co-regulators that are overexpressed in prostate cancer [10–12]. Targeting these HATs reduces prostate cancer cell growth [13]. BRD4, a chromatin reader that recognizes acetylated histones, is another AR co-regulator that is upregulated in CRPC [14]. Inhibition of BRD4 via a BET inhibitor reduces its interaction with AR, inhibits AR transcriptional activity, and reduces prostate cancer cell growth *in vitro* and *in vivo* [15, 16]. Furthermore, BET inhibitors have advanced to clinical trials for CRPC [17].

AR co-regulators implicated in prostate cancer progression also include transcription factors, such as members of the ETS domain family of transcription factors that recognize ETS binding motifs in the genome [7, 18]. ETS binding motifs are enriched at AR occupancy sites in prostate cancer cells, and multiple ETS family members are upregulated in prostate cancer [18–22]. In particular, the ETS family member ELK1 was found to interact with the N-terminal domain (NTD) of AR and directly regulate its recruitment to chromatin in prostate cancer cells [23, 24]. Inhibition of ELK1 reduces expression of AR target genes and suppresses prostate cancer cell growth [23, 24].

However, of the hundreds of proteins that have been identified as AR co-regulators, only a small portion have been definitively linked to disease; furthermore, AR co-regulators function in multi-protein transcriptional complexes [7–9, 25]. Therefore, it is important to identify and characterize co-regulators that play a key role in mediating these complexes, are crucial for driving AR transcriptional activity and growth, and are upregulated in prostate cancer.

To this end, our lab previously performed an unbiased, genome-wide siRNA screen for novel AR co-regulators, and identified MED19, a subunit of the middle module of the Mediator complex that functionally bridges promoters and enhancers to connect transcription factors and RNA polymerase II (Pol II) [26, 27]. We found that MED19 depletion greatly inhibited AR transcriptional activity and proliferation of LNCaP-abl cells (androgen-independent) and LNCaP cells (androgen-dependent). MED19 mRNA is also upregulated in primary and metastatic prostate cancer, and its abundance correlates with lower overall survival [26, 28, 29]. From this, we proposed that upregulation of MED19 in prostate cancer cells drives AR activity and androgen independence. However, the mechanism by which MED19 regulates AR activity, particularly under low androgen conditions (as occurs during ADT), as well as the downstream gene targets controlling growth, remained unknown.

In this study we examined the ability of MED19 to confer androgen independence and its effect on gene expression and AR occupancy. We identified the specific gene target, MAOA, and cooperating transcription factor, ELK1, underlying MED19 regulation of androgen-independent growth.

## Results

### MED19 overexpression promotes androgen independence and confers a growth advantage to prostate cells

To determine whether MED19 is sufficient to convert androgen-dependent prostate cancer cells to androgen independence, we stably overexpressed MED19 in the prototypical androgen-dependent LNCaP cell line (MED19 LNCaP cells) (S1A and S1B Fig). As a control, we also created the parental LNCaP cells stably expressing the empty lentiviral vector (control LNCaP cells). Both lines represent a pool of cells. We compared their proliferation under androgen-deprived conditions in 2D culture; by colony formation, which measures the survival and replicative potential of individual cancer cells; and by spheroid formation, a 3D cell culture condition more representative of a tumor.

In contrast to control LNCaP cells, MED19 LNCaP cells showed robust proliferation in 2D culture, increased colony formation, and larger spheroid formation, when cultured in media depleted of steroids (Fig 1A-1C). This demonstrates that expression of MED19 is sufficient to promote androgen independence in prostate cancer cells. This is consistent with our previous findings that MED19 in androgen-independent LNCaP-abl cells is necessary for growth. MED19 overexpression also conferred a growth advantage, albeit less striking, when the cells were cultured in complete media containing endogenous steroids (Fig 1D-1F). This is also consistent with reports from our lab and others demonstrating that depletion of MED19 inhibits LNCaP cell growth in the presence of androgens [26, 30].

**Fig 1.**
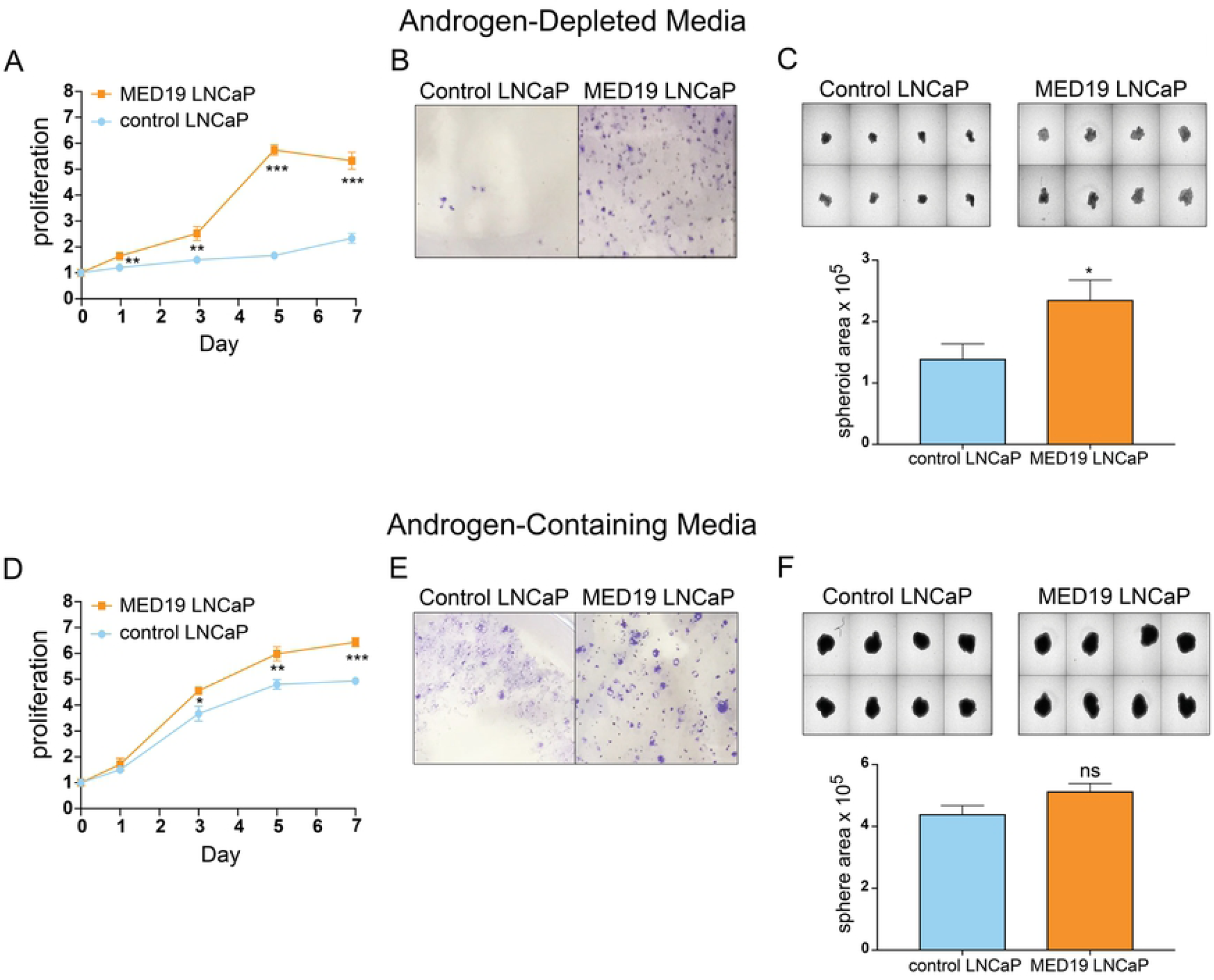
MED19 overexpression confers a growth advantage and enables androgen-independent growth of LNCaP cells. LNCaP cells stably overexpressing MED19 (MED19 LNCaP cells) and LNCaP cells expressing the empty lentiviral vector (control LNCaP cells) were cultured in media depleted of androgens by addition of FBS charcoal-stripped of steroids (A, B, C) or media containing androgens by addition of standard FBS (D, E, F). A) and D) Proliferation was measured over 7 days and is expressed as fold change in relative fluorescent units (RFU), normalized to Day 0. B) and E) Colony formation was evaluated by culturing MED19 LNCaP cells and control LNCaP cells at low density for 11 days and fixing and staining with crystal violet. C) and F) Spheroid formation was evaluated by culturing cells on low attachment plates for 10 days and quantifying average spheroid area. Experiments were performed in biological triplicate, with representative results shown. *p < 0.05; **p < 0.01; and ***p < 0.001.

We also examined if upregulation of MED19 could promote the proliferation of other early stage prostate cancer cell lines. Indeed, we found that overexpression of MED19 increased proliferation (as well as colony formation) in non-malignant RWPE-1 cells (Fig 2A and S2A Fig), but did not affect the growth of RWPE-2 cells, which are a malignant derivative of RWPE-1 cells, transformed with RAS (Fig 2B and S2B Fig) [31]. This is in spite of similar levels of MED19 protein expressed in RWPE-1 cells and RWPE-2 cells (S3 Fig). Furthermore, a murine prostate stem cell line transformed with activated AKT grew markedly faster upon MED19 overexpression compared to its control counterpart (Fig 2C and S2C Fig). This was recapitulated in a xenograft model, where the MED19-overexpressing cells produced larger tumors than controls (Fig 2D). This corroborates the growth advantage of MED19 found in LNCaP cells. Together, this indicates a role for MED19 in conversion of early stage cells to aggressive growth and androgen independence.

**Fig 2.**
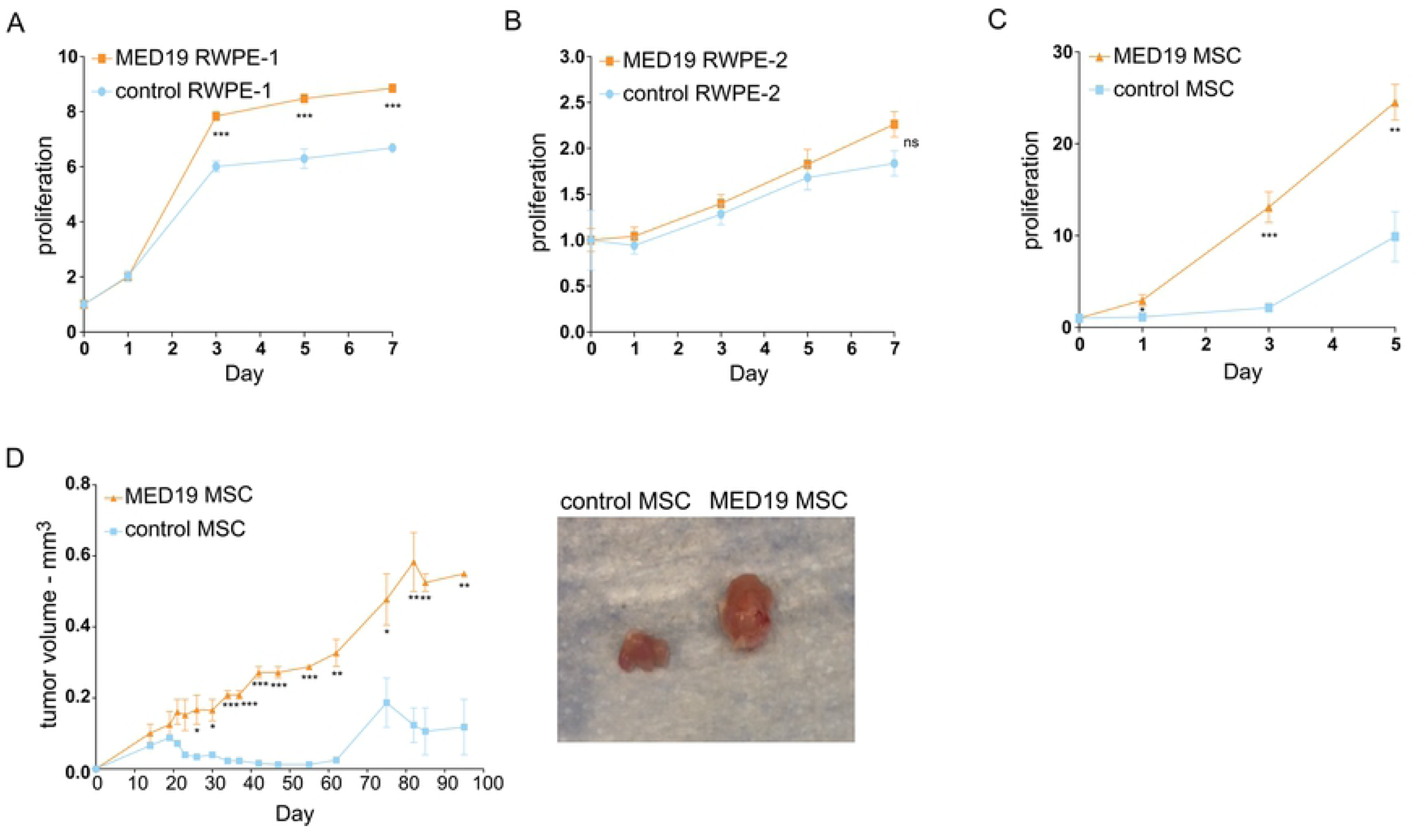
MED19 overexpression promotes growth *in vitro* in non-malignant RWPE-1 cells and *in vitro* and *in vivo* in mouse prostate stem cells. A) RWPE-1, B) RWPE-2, or C) mouse stem cells with activated AKT (MSC), stably overexpressing MED19 (MED19 RWPE-1/RWPE-2/MSC) or control empty vector (control RWPE-1/RWPE-2/MSC), were cultured in their standard media. A-C) Proliferation was measured over 5 days for the MSC, which have a rapid doubling time, or 7 days for RWPE-1 cells and RWPE-2 cells, and is expressed as fold change in relative fluorescent units (RFU) normalized to Day 0. Experiments were performed in biological duplicate, with representative results shown. D) MED19 MSC or control MSC were injected into the flanks of Nu/J mice and tumor volume was measured over 95 days (2 mice per group). Representative images of tumors taken at time of sacrifice are shown. *p < 0.05; **p < 0.01; and ***p < 0.001.

### MED19 depends on AR activity for its growth advantage but does not alter AR expression

As AR amplification is a common mechanism to achieve androgen independence, we evaluated the mRNA and protein levels of AR with MED19 overexpression in LNCaP cells. We found that AR mRNA and protein were unchanged in MED19 LNCaP cells compared to control LNCaP cells under androgen deprivation (Fig 3A and 3B). MED19 overexpression also did not induce expression of AR-V7, a constitutively active splice variant of AR lacking the ligand binding domain that can drive androgen independence in prostate cancer (S4 Fig) [32].

**Fig 3.**
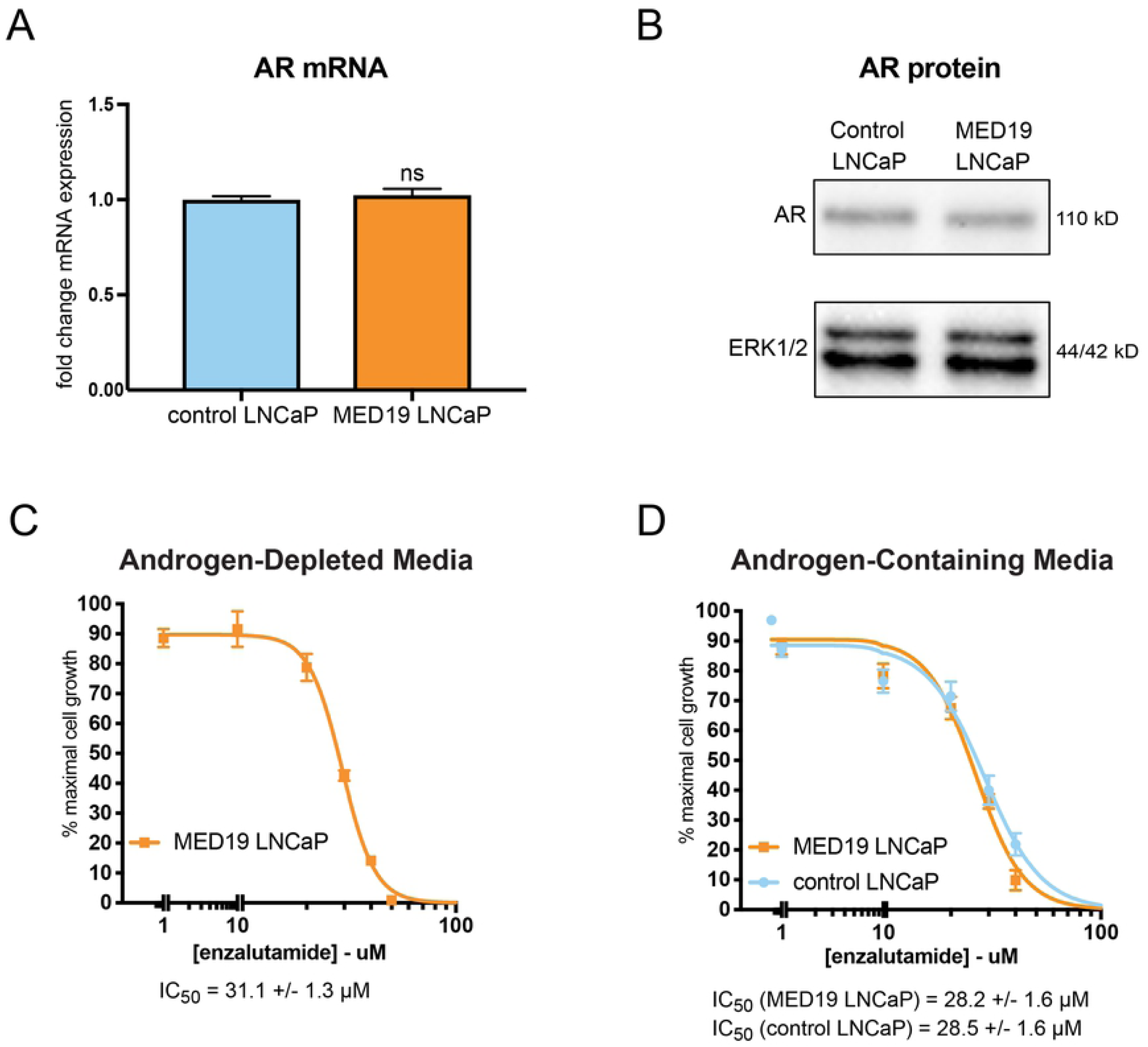
MED19 LNCaP cells depend on AR transcriptional activity for androgen-independent growth and do not have altered expression of AR. A) RNA was extracted from MED19 LNCaP cells and control LNCaP cells cultured under androgen deprivation for 3 days, and AR mRNA measured by qPCR (fold change mRNA expression normalized to RPL19, with AR mRNA expression in control LNCaP cells set as “1”). B) Total protein lysate was collected and probed for AR protein levels by western blot. ERK1/2 was used as a loading control. C) and D) MED19 LNCaP cells were treated with enzalutamide (0-80 µM) in C) androgen-depleted media, or D) androgen-containing media, alongside control LNCaP cells. Proliferation was measured over 7 days. Percent cell growth at day 7 is normalized to vehicle treatment (0 µM, 100%). The IC_50_ from three experiments is shown. Experiments were performed in biological triplicate, with representative results shown. *p < 0.05; **p < 0.01; and ***p < 0.001.

We then examined the reliance of MED19 LNCaP cells on AR for their androgen-independent growth by determining their sensitivity to enzalutamide, an AR antagonist that reduces AR transcriptional activity in part by preventing AR nuclear accumulation [33]. Enzalutamide inhibited the proliferation of MED19 LNCaP cells both in the presence and absence of androgens, indicating that the growth advantage conferred by MED19 requires AR transcriptional activity (Fig 3C and 3D). We confirmed these results by siRNA depletion of AR, which also reduced androgen-dependent and androgen-independent growth of MED19 LNCaP cells (S5A and S5B Fig).

### MED19 regulates gene expression by altering AR occupancy and H3K27 acetylation at target genes

MED19 relies on the transcriptional activity of AR for its growth advantage, and as part of the Mediator complex controls gene expression through transcription factor, co-regulator, and histone modifying complex recruitment. Thus, we evaluated the effect of overexpressed MED19 on gene expression, AR occupancy, and H3K27 acetylation under androgen deprivation and in response to androgens, with a particular focus to identify the specific gene expression and AR occupancy changes driving androgen independence. To this end, we performed RNA sequencing (RNA-seq) and ChIP sequencing (ChIP-seq) studies for FLAG-tagged MED19, AR, and H3K27 acetylation in MED19 LNCaP cells and control LNCaP cells cultured under androgen deprivation and with treatment of R1881, a synthetic AR agonist.

Global AR occupancy under androgen deprivation, as measured by total number of and individual level of occupancy at AR sites in published ChIP-seq studies, is low compared to androgen treatment. We speculated that MED19 may alter AR activity under androgen deprivation by modulating AR at low occupancy sites. Therefore, we included in our ChIP-seq study all AR-specific sites, including those with low occupancy in androgen-deprived conditions. We used rigorous quality controls to maximize capture of AR sites and ensure strict specificity to AR occupancy (see detailed ChIP-seq section under Materials and Methods) (S6C Fig and S7 Table).

Under androgen deprivation, there was a striking and very selective change in gene expression profile with MED19 overexpression, with a total of 151 genes altered (76 genes upregulated and 75 genes downregulated, fold change≥1.5 and p-adj≤0.05) (Fig 4A and S1 Table). This was accompanied by a selective shift in the AR cistrome (∼12% of total AR sites are occupied only in control LNCaP cells or only in MED19 LNCaP cells), without a global change in the total number of sites occupied by AR (Fig 4A). As expected, with androgen treatment, the total number of AR sites increased (Figs 5A and 5B). There was a selective shift in the AR cistrome when MED19 is overexpressed in the presence of androgens as well (Fig 4B). There was also a shift in gene expression: 309 genes were differentially expressed with MED19 overexpression in the presence of androgens (78 genes upregulated and 231 downregulated, fold change≥1.5 and p-adj≤0.05) (Fig 4B and S2 Table).

**Fig 4.**
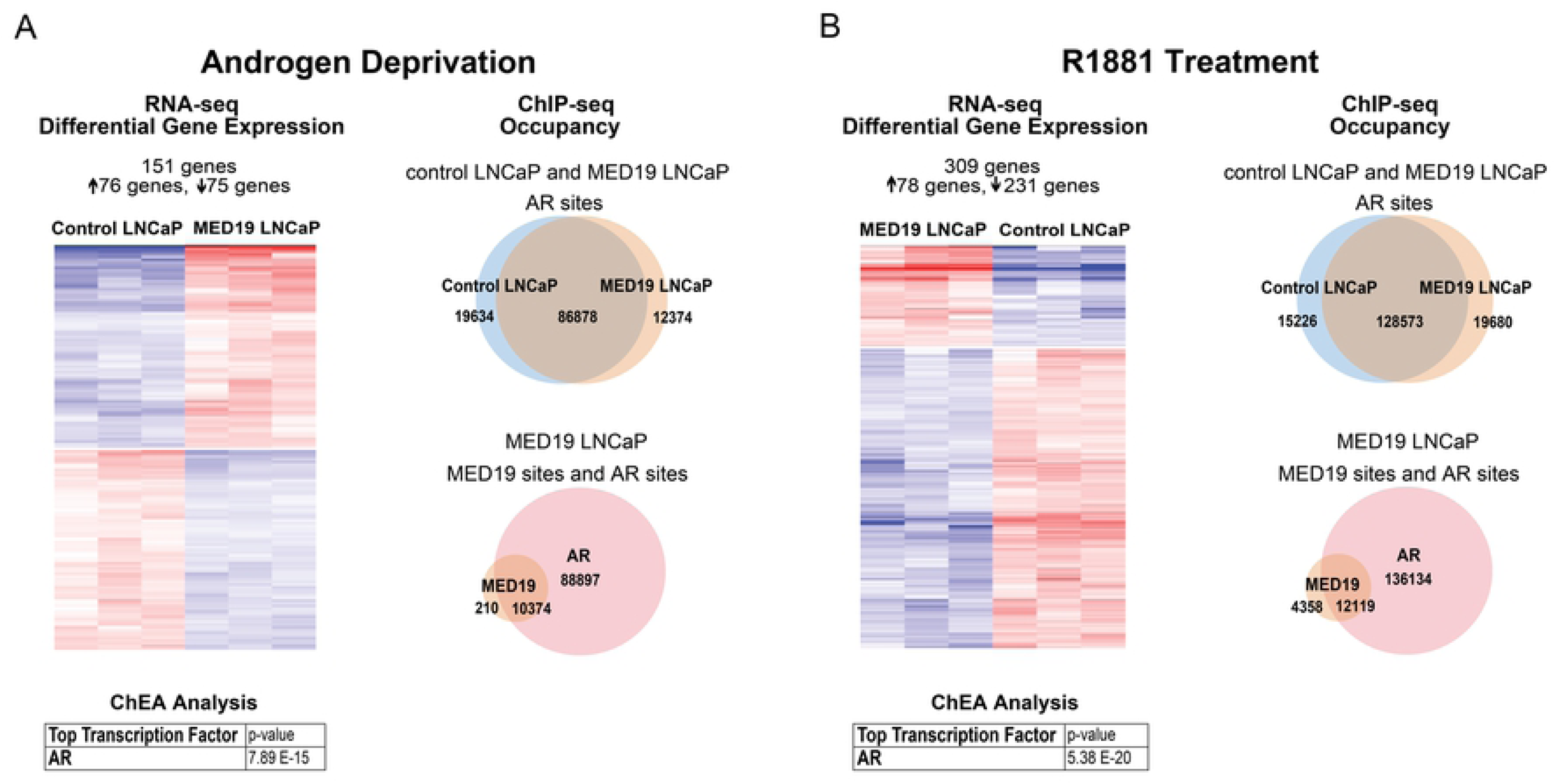
MED19 overexpression causes a selective shift in gene expression and in the AR cistrome under androgen deprivation and with R1881 treatment. MED19 LNCaP cells and control LNCaP cells were cultured under androgen deprivation for 3 days, and cells were treated with ethanol vehicle or 10 nM R1881 overnight (16 h). RNA-seq with ribodepletion was performed in biological triplicate. ChIP-seq for FLAG-MED19, AR, and H3K27ac was performed in biological triplicate. A) (Left) Heatmap of differentially expressed genes with MED19 overexpression (fold change ≥1.5, p-adj ≤0.05) for androgen deprivation, associated with AR as top regulatory transcription factor from ChEA. (Right) Number and overlap of occupancy sites under androgen deprivation for AR in control LNCaP cells and MED19 LNCaP cells (top) and for AR and MED19 in MED19 LNCaP cells (bottom). B) (Left) Heatmap of differentially expressed genes with MED19 overexpression (fold change ≥1.5, p-adj ≤0.05) for R1881 treatment, associated with AR as top regulatory transcription factor from ChEA. (Right) Number and overlap of occupancy sites with R1881 treatment for AR in control LNCaP cells and MED19 LNCaP cells (top) and for AR and MED19 in MED19 LNCaP cells (bottom).

Of these 309 genes, 82 were also differentially expressed in the absence of androgens (∼50% of total genes differentially expressed in the absence of androgens) (S1 and S2 Tables). This holds true for MED19 occupancy as well: the total number of MED19 sites increased with androgen treatment, with ∼50% of the sites occupied in the absence of androgens also occupied in the presence of androgens (Fig 5B). This indicates that there is partial overlap in MED19 regulation of gene expression and AR activity in the absence and presence of androgens, consistent with the differential growth advantage in the absence and presence of androgens when MED19 is overexpressed.

**Fig 5.**
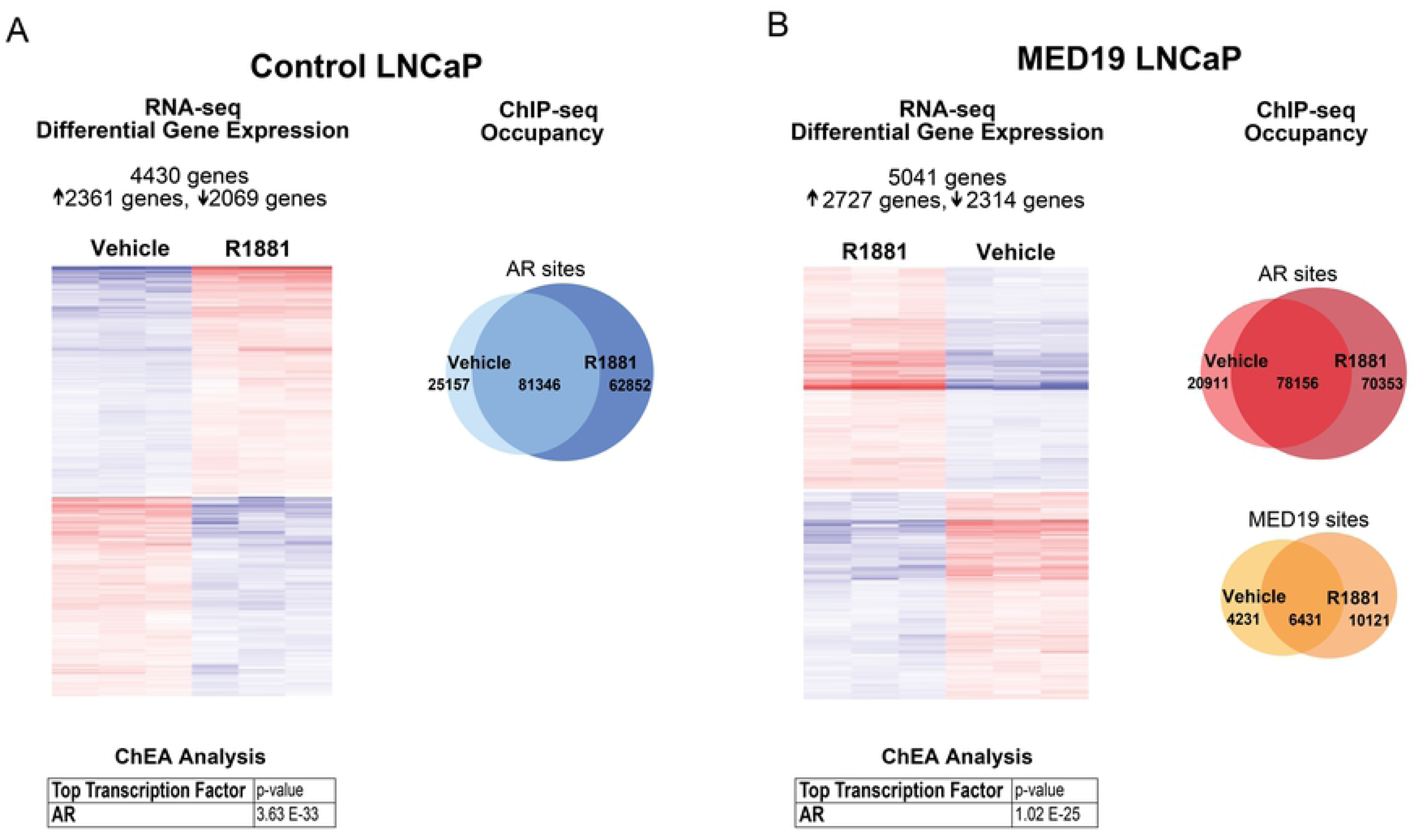
MED19 overexpression alters the response to androgens. MED19 LNCaP cells and control LNCaP cells were cultured under androgen deprivation for 3 days, and cells were treated with ethanol vehicle or 10 nM R1881 overnight (16 h). RNA-seq with ribodepletion was performed in biological triplicate. ChIP-seq for FLAG-MED19, AR, and H3K27ac was performed in biological triplicate. A) (Left) Heatmap of differentially expressed genes (fold change ≥1.5, p-adj ≤0.05) for control LNCaP cells treated with R1881 vs. vehicle, associated with AR as top regulatory transcription factor from ChEA. (Right) Number and overlap of occupancy sites for R1881 vs. vehicle treatment for AR in control LNCaP cells. B) (Left) Heatmap of differentially expressed genes (fold change ≥1.5, p-adj ≤0.05) for MED19 LNCaP cells treated with R1881 vs. vehicle, associated with AR as top regulatory transcription factor from ChEA. (Right) Number and overlap of occupancy sites for R1881 vs. vehicle treatment for AR (top) and for MED19 (bottom) in MED19 LNCaP cells.

MED19 occupancy in the absence and presence of androgens corresponds almost entirely with AR occupancy, with virtually every gene differentially expressed in MED19 LNCaP cells occupied by AR, and many (the majority in the absence of androgens) also occupied by MED19, indicating direct regulation by MED19 (Fig 4A and 4B, S6 Table). In fact, most of the MED19-regulated genes are androgen-responsive, and many have been reported as AR target genes (S1 and S2 Tables). AR was also the top predicted regulatory transcription factor candidate using Chromatin immunoprecipitation Enrichment Analysis (ChEA) (Fig 4A and 4B, S1 and S2 Tables). This confirms that MED19 regulation of gene expression is driven by AR.

In response to androgen treatment, ∼4500 genes in control LNCaP cells and ∼5000 genes in MED19 LNCaP cells were differentially expressed (≥1.5-fold, p-adj≤0.05), and, as expected, AR was the top transcription factor from ChEA analysis for both (Fig 5A and 5B, S3 and S4 Tables). This included expected changes in canonical AR target genes, such as upregulation of *FKBP5* and *PSA* (S1 and S2 Tables, S7A and S7B Fig). Some genes were differentially expressed in response to androgens unique to control LNCaP cells (645 genes) or to MED19 LNCaP cells (1250 genes), comprising ∼15 or ∼25% of the total genes differentially expressed in response to androgens in control LNCaP cells or MED19 LNCaP cells, respectively, indicating that MED19 alters which genes AR regulates in response to androgens (S5 Table).

However, MED19 appears mainly to modulate the response of canonically androgen-regulated genes: the top 100 androgen-induced and androgen-repressed genes almost all overlapped between MED19 LNCaP cells and control LNCaP cells, with MED19 overexpression augmenting the response to androgen for some genes, and reducing the response to androgen for others (S5 Table). However, the overall response to androgens does not appear to markedly differ with MED19 overexpression, and differential gene expression with MED19 overexpression in the presence and absence of androgens is very selective (Figs 4 and 5). This indicates that MED19 does not alter the entire AR-regulated transcriptome, nor the global response to androgens (Figs 4 and 5). Overall, this suggests that MED19 alters the cellular response to androgens in a specific manner, consistent with the growth advantage conferred by MED19 overexpression in the presence of androgens.

Although AR occupies unique sites in MED19 LNCaP cells, the majority of genes altered with MED19 overexpression contain AR sites shared by control LNCaP cells and MED19 LNCaP cells (S6 Table). What we observed at a number of these sites was a change in the level of AR occupancy and/or H3K27 acetylation beyond a “present/absent” or “on/off” binary. These subtle changes in gene occupancy corresponded with MED19 activation or repression of genes from the RNA-seq study, indicating that MED19 alters gene expression through small shifts in AR occupancy and H3K27 acetylation. We also observed that the changes in AR occupancy and H3K27 acetylation, like the changes in gene expression with MED19 overexpression, did not simply mimic the changes that occurred with androgen treatment.

For example, LRRTM3 (Leucine Rich Repeat Transmembrane Neuronal 3) is one of the most upregulated genes upon MED19 overexpression under androgen deprivation, while androgen treatment suppresses LRRTM3 expression (S1-S4 Tables, Fig 6A). With MED19 overexpression, there is a clear increase in AR occupancy and H3K27 acetylation at several regulatory intronic sites at LRRTM3, with MED19 occupancy at one of these sites (Fig 6B, S6 Table). Conversely, androgen treatment reduces H3K27 acetylation and alters AR occupancy (S8A and S8B Fig and S6 Table). In contrast, MAST4 is one of the most downregulated genes with MED19 overexpression under androgen deprivation, and is also suppressed by androgen treatment (S1-S4 Tables, S9A Fig). There is a clear reduction in H3K27 acetylation adjacent to the MAST4 promoter with MED19 overexpression and with androgen treatment. MED19 overexpression and androgen treatment induce a reorganization of AR occupancy (including a site of occupancy with MED19), though the former does not exactly mimic the latter (S9B Fig, S6 Table). Thus, it appears that AR occupancy at specific targets is altered by MED19 overexpression. This may be responsible for the changes in gene expression and attendant effects on cell proliferation.

**Fig 6.**
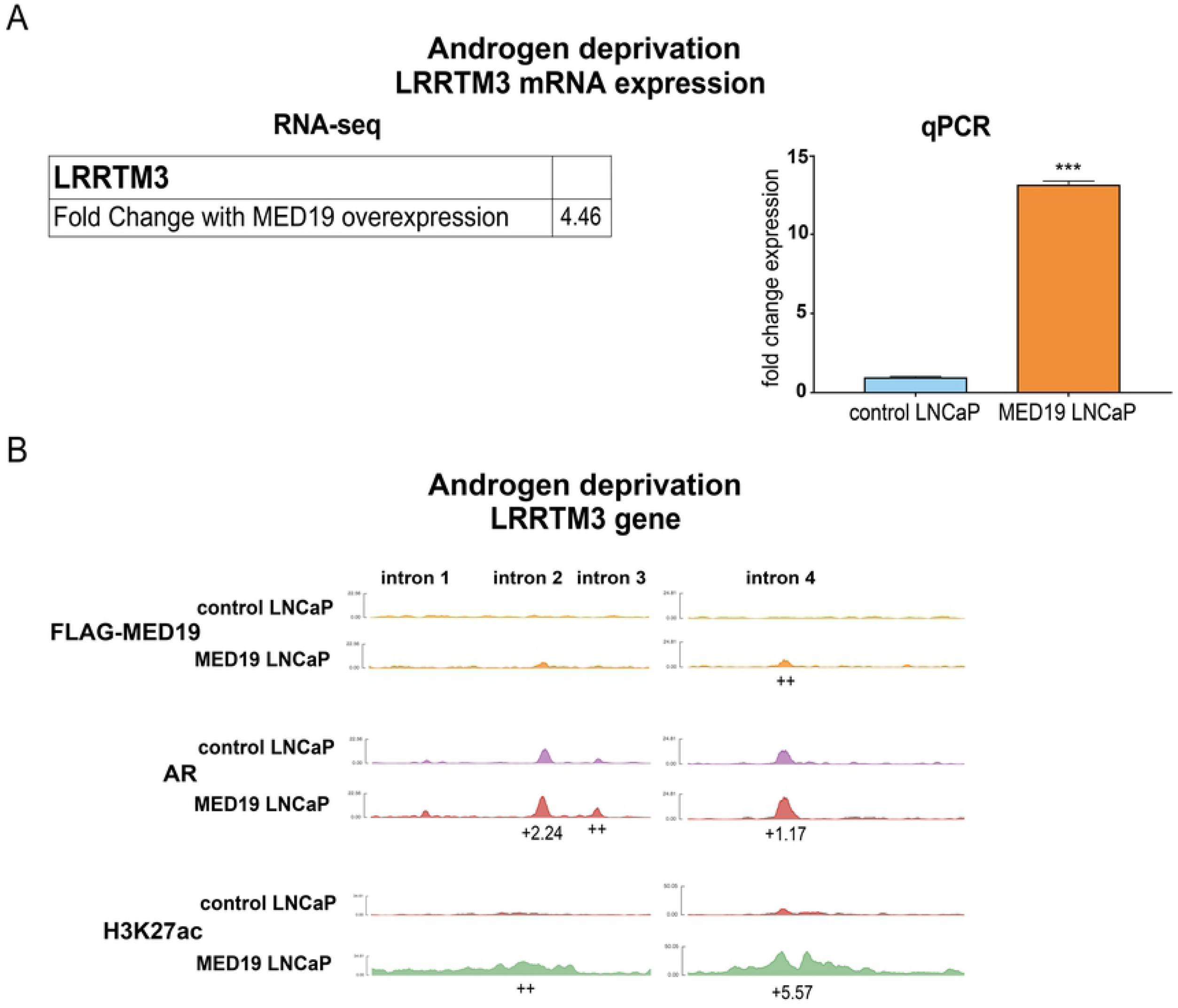
MED19 occupies gene targets like LRRTM3 under androgen deprivation and alters mRNA expression, AR occupancy, and H3K27 acetylation. MED19 LNCaP cells and control LNCaP cells were cultured under androgen deprivation for 3 days and treated overnight (16 h) with ethanol vehicle (shown) or 10 nM R1881 (shown in S8 Fig). ChIP-seq for FLAG-MED19, AR, and H3K27ac was performed in biological triplicate. A) Fold change mRNA expression from RNA-seq, and qPCR validation of upregulation of LRRTM3 mRNA expression under androgen deprivation (performed in biological triplicate with representative result shown; fold change expression normalized to RPL19 with LRRTM3 mRNA expression in control LNCaP cells set as “1”). *p < 0.05; **p < 0.01; and ***p < 0.001. B) ChIP-seq tracks (representative results) for FLAG-MED19, AR, and H3K27ac under androgen deprivation shown for intronic regions of LRRTM3. Fold change (up (+) or down (-)) in occupancy scores for MED19 LNCaP cells compared to control LNCaP cells shown for each peak (see Table S6 for all occupancy scores). ++ indicates positive occupancy score in MED19 LNCaP cells and a score of zero in control LNCaP cells; -- indicates an occupancy score of zero in MED19 LNCaP cells and a positive score in control LNCaP cells.

We wanted to determine the specific gene targets altered by MED19 overexpression that were responsible for promoting androgen-independent growth. We decided to focus on genes upregulated by MED19 overexpression under androgen deprivation, which could be depleted to inhibit androgen-independent growth. Given that LRRTM3 appears to be a direct target of MED19, with changes in AR occupancy that correlated with a large upregulation in expression, we tested its effect on proliferation. However, LRRMT3 depletion had very little effect on androgen-independent growth (S8C Fig). Furthermore, LRRTM3 has no published connection to AR or prostate cancer. Therefore, we stratified for gene targets occupied by MED19 and AR with an established connection to AR, preferably AR target genes, and known to play a role in prostate cancer proliferation.

### MED19 upregulates expression and promotes AR occupancy and H3K27 acetylation at MAOA, which is required for androgen-independent growth

One target that fulfilled these criteria is MAOA (monoamine oxidase A). MAOA is a mitochondrial enzyme that degrades monoamine neurotransmitters and dietary amines and produces hydrogen peroxide. It has a well-established role in promoting aggressive prostate cancer cell growth, invasion, and metastasis [34–37]. MAOA is also reported as an AR target gene with an androgen response element (ARE) in its promoter [38]. Indeed, MAOA expression increased in control LNCaP cells in response to R1881, which was comparable to the increase in expression in MED19 LNCaP cells under androgen deprivation (Fig 7A, S1 and S3 Tables, S10A Fig). This indicates that for MAOA expression, MED19 overexpression under androgen deprivation recapitulates the effects of androgen activation.

**Fig 7.**
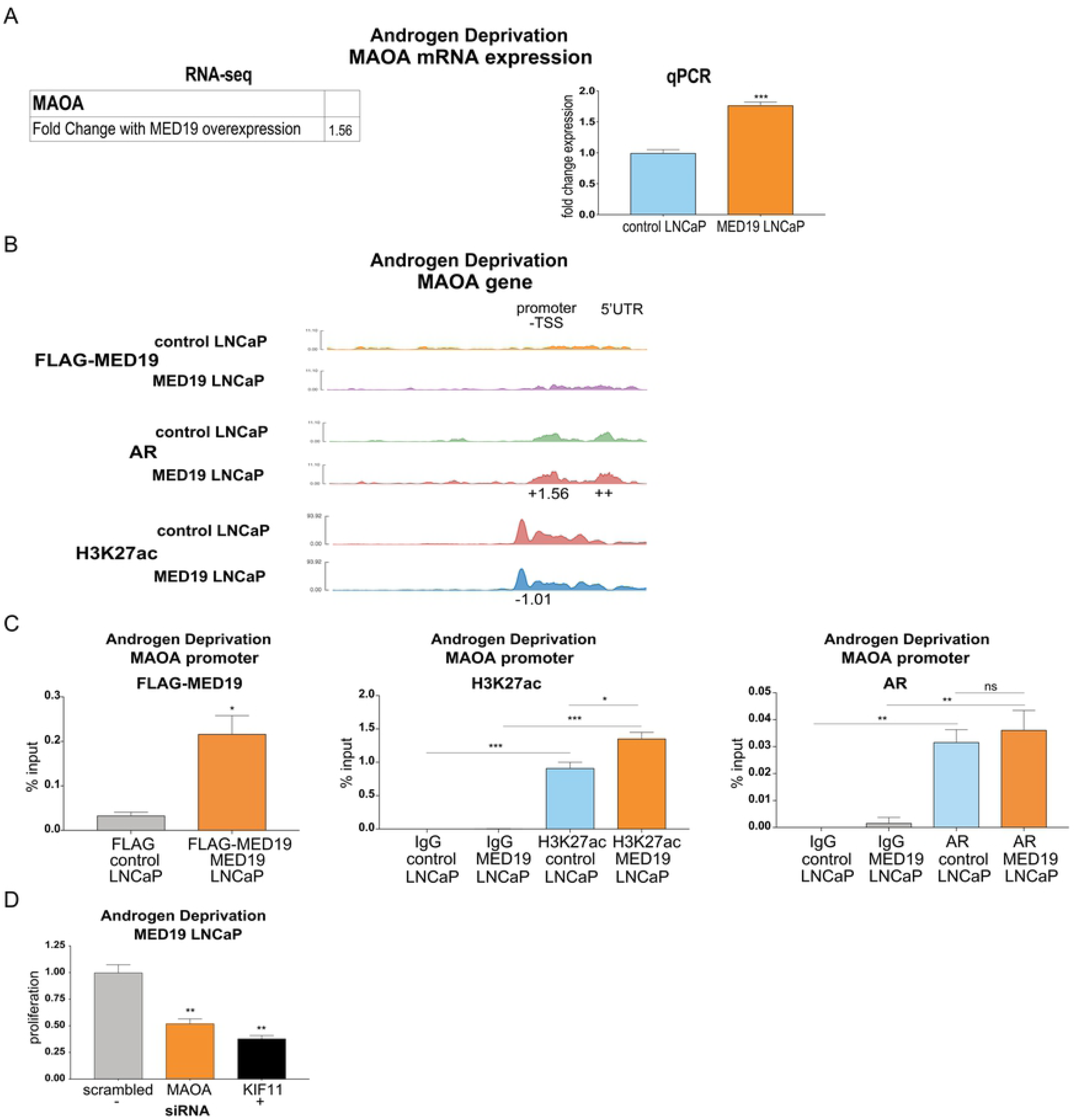
MED19 promotes AR occupancy and H3K27 acetylation at MAOA under androgen deprivation, which controls androgen-independent growth. MED19 LNCaP cells and control LNCaP cells were cultured under androgen deprivation for 3 days and treated overnight (16 h) with ethanol vehicle (shown) or 10 nM R1881 (shown in S10 Fig). ChIP-seq for FLAG-MED19, AR, and H3K27ac was performed in biological triplicate. A) Fold change mRNA expression from RNA-seq, and qPCR validation of upregulation of MAOA mRNA expression under androgen deprivation (performed in biological triplicate with representative result shown; fold change expression normalized to RPL19 with MAOA mRNA expression in control LNCaP cells set as “1”). B) ChIP-seq tracks (representative results) for FLAG-MED19, AR, and H3K27ac under androgen deprivation shown for promoter region of MAOA. Fold change (up (+) or down (-)) in occupancy scores for MED19 LNCaP cells compared to control LNCaP cells shown for each peak (see Table 6 for all occupancy scores). ++ indicates positive occupancy score in MED19 LNCaP cells and a score of zero in control LNCaP cells; -- indicates an occupancy score of zero in MED19 LNCaP cells and a positive score in control LNCaP cells. C) ChIP-qPCR for FLAG-MED19, AR, and H3K27ac at MAOA promoter overlapping with published ARE. Experiment was performed in duplicate, with representative results shown. D) MAOA was depleted by siRNA and proliferation of MED19 LNCaP cells in androgen-depleted media was evaluated after 7 days, normalized to proliferation with scrambled siRNA treatment (negative control, light grey). KIF11 knockdown is included as a positive control (black). Experiment was performed in biological duplicate, with representative results shown. *p < 0.05; **p < 0.01; and ***p < 0.001.

Indeed, AR occupies the promoter and 5’ UTR of MAOA, with increased and reorganized occupancy when MED19 is overexpressed and when the cells are treated with R1881 (Fig 7B, S10B Fig). ChIP-qPCR for the MAOA promoter region overlapping with the published ARE confirmed MED19 and AR occupancy, as well as increased H3K27 acetylation with MED19 overexpression and with R1881 treatment (Fig 7C, S10C Fig). To determine if MED19 activation of MAOA was responsible for androgen-independent growth, we depleted MAOA in MED19 LNCaP cells under androgen deprivation and measured proliferation. MAOA depletion reduced growth by ∼50% (Fig 7D). Interestingly, from the RNA-seq study, MAOA is not differentially upregulated with MED19 overexpression in the presence of androgens, which is consistent with the relatively smaller growth advantage of MED19 overexpression when androgens are present (S1-S4 Tables, S10A Fig).

### ELK1 is enriched at MED19 and AR occupied sites and upregulated targets, driving MAOA expression and androgen-independent growth

Given that increased or decreased AR recruitment corresponded to activation or repression of target genes as a function of MED19 overexpression, and given that AR works in concert with other transcription factors to control gene expression, we determined the identity of other transcription factor binding motifs associated with AR and MED19 occupancy. We were particularly interested in identifying any transcription factors uniquely enriched in MED19 LNCaP cells under androgen deprivation, that correlated with MED19 occupancy, and would have an established connection to prostate cancer and regulation of MAOA.

FOXA1 and FOXM1 were the most enriched transcription factor motifs for AR occupancy in control LNCaP cells and MED19 LNCaP cells, in the absence or presence of androgens, as well as for MED19 occupancy in the absence or presence of androgens (S11A and S11B Fig). This is consistent with the well-established role of FOXA1 as a major AR co-regulator and pioneer factor in prostate cancer cells, and the emerging role of FOXM1 in this capacity as well [39–41]. This is also consistent with the RNA-seq analysis and the vast majority of MED19 sites overlapping with AR sites. However, FOXA1- and FOXM1-mediated recruitment of AR seems unlikely to be the dominant mechanism by which overexpressed MED19 promotes gene expression changes, given that FOXA1 and FOXM1 sites are highly enriched in both MED19 LNCaP cells and control LNCaP cells in all conditions (S11A and S11B Fig).

We then focused on the intersection between MED19 and AR occupancy at sites engaged by AR *uniquely* in MED19 LNCaP cells. Under androgen deprivation, we found that the most enriched motif corresponded to ELK1 (Fig 8A). ELK1 is an ETS transcription factor and AR co-regulator that promotes growth in prostate cancer cells and regulates ligand-independent recruitment of AR to chromatin through interaction with the AR NTD [23, 24]. Interestingly, also enriched under androgen deprivation, as well as with androgen treatment, were several other members of the ETS family of transcription factors (Fig 8A and S12A Fig). In the presence of androgens, the most enriched motif corresponded not to ELK1 but to SP1, an AR-interacting protein upregulated in prostate cancer (S12A Fig) [42]. SP1 is reported to promote AR target gene expression in response to androgens and to occupy sites near gene promoters [43, 44]. This confirms that MED19 likely regulates AR occupancy and activity at its upregulated targets through different mechanisms in the absence and presence of androgens.

**Fig 8.**
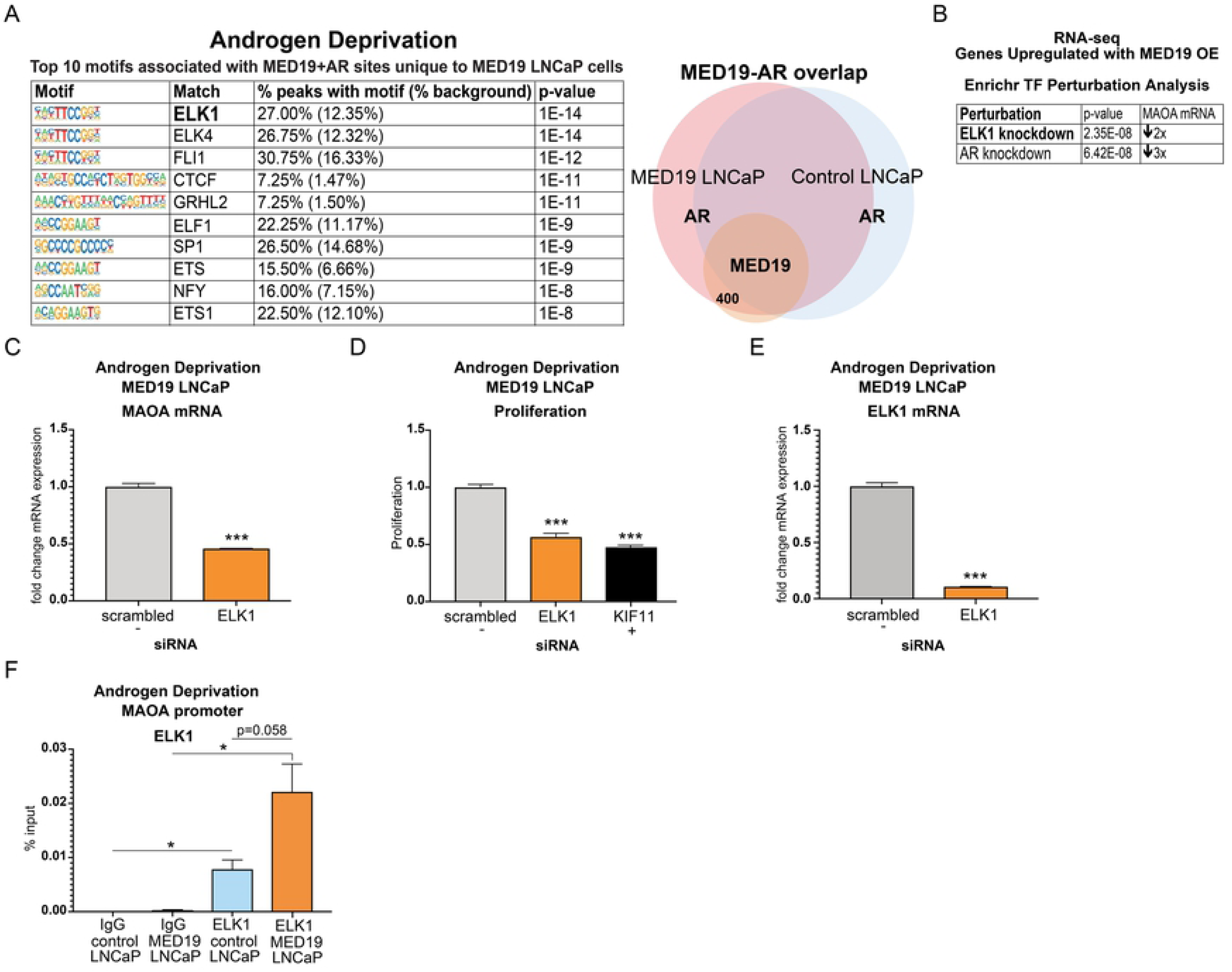
ELK1 is enriched at sites of AR and MED19 occupancy unique to MED19 LNCaP cells under deprivation, occupies the MAOA promoter, and controls MAOA expression and androgen-independent growth. A) MED19 LNCaP cells and control LNCaP cells were cultured under androgen deprivation for 3 days and treated overnight with ethanol vehicle. ChIP-seq for FLAG-MED19, AR, and H3K27ac was performed in biological triplicate. The top 10 enriched transcription factor motifs under androgen deprivation associated with sites of AR and MED19 occupancy in MED19 LNCaP cells where AR is present only in MED19 LNCaP cells are shown, with ELK1 as the top associated transcription factor (labeled diagram of occupancy sites, right). B) ELK1 knockdown is the top hit from Transcription Factor Perturbation from Enrichr associated with genes upregulated by MED19 overexpression under androgen deprivation from the RNA-seq study; AR knockdown is also strongly associated. Fold downregulation of MAOA mRNA with associated ELK1 knockdown (GSE 34589) or associated AR knockdown (GSE 22483). C) ELK1 was depleted by siRNA and mRNA expression of MAOA in MED19 LNCaP cells cultured in androgen-depleted media was measured after 5 days (fold change expression normalized to RPL19, with MAOA mRNA expression with scrambled siRNA treatment set as “1”). Experiment was performed in biological triplicate, with representative results shown. D) ELK1 was depleted by siRNA and proliferation of MED19 LNCaP cells in androgen-depleted media was evaluated after 7 days, normalized to proliferation with scrambled siRNA (negative control, light grey). KIF11 knockdown is included as a positive control (black). Experiment was performed in biological duplicate, with representative results shown. E) ChIP-qPCR for ELK1 at the MAOA promoter overlapping with published ARE. *p < 0.05; **p < 0.01; and ***p < 0.001.

Interestingly, although sites of AR (and MED19) occupancy in MED19 LNCaP cells and in control LNCaP cells, in the presence and absence of androgens, were enriched for ARE and other canonical AR-related motifs (i.e. ARE half-site and FOXA1:AR motifs), as expected, this enrichment was reduced under androgen deprivation (S13A-S13C Fig). In addition, these motifs were absent at sites of MED19 and AR occupancy in MED19 LNCaP cells where AR was present only in MED19 LNCaP cells. This would be consistent with the ability of ELK1 to act as an AR tethering protein to promote AR recruitment to non-canonical sites.

We next used Enrichr’s “Transcription Factor Perturbation” tool to compare published gene expression changes with transcription factor knockdown or overexpression to our RNA-seq study. Strikingly, under androgen deprivation, genes upregulated with MED19 overexpression corresponded consistently to genes downregulated with ELK1 knockdown (top hit) or with AR knockdown, including MAOA (Fig 8B, S12B Fig). However, with R1881 treatment, SP1 was not associated with genes upregulated with MED19 overexpression in the presence of androgens; rather, these were associated with SRF, which was not enriched in the ChIP-seq data (S12C Fig). Given the strong connection to ELK1, we tested its functional role in gene expression and androgen-independent growth. We depleted ELK1 by siRNA in MED19 LNCaP cells and measured the effect on MAOA mRNA expression and proliferation under androgen deprivation. ELK1 knockdown both greatly reduced expression of MAOA and inhibited androgen-independent growth (Fig 8C-8E).

To determine if ELK1 occupied MAOA at sites where AR and MED19 were present, we performed ChIP-qPCR for ELK1 adjacent to the MAOA promoter, overlapping with the reported ARE, where AR, MED19, and H3K27 acetylation were present (Fig 8F, S14 Fig). Indeed, we found that ELK1 also occupied this region, with a trend toward increased ELK1 occupancy under androgen deprivation when MED19 is overexpressed (Fig 8F). This indicates that MED19 and ELK1 cooperate to further AR occupancy and H3K27 acetylation, increase MAOA expression, and promote androgen-independent growth.

## Discussion

We have demonstrated that overexpression of MED19 in androgen-dependent LNCaP cells provides a growth advantage in the absence and presence of androgens. This is mediated by AR. The cells remain dependent on AR for growth under androgen deprivation, without increasing full-length AR abundance or splice variant AR-V7 expression. Therefore, increased expression of MED19 is sufficient to convert a cell that is androgen-dependent to one that is androgen-independent for growth.

Consistent with this are reports from our lab and others that MED19 depletion reduced AR transcriptional activity and growth of LNCaP cells and LNCaP-abl cells, which are derived from LNCaP cells and are androgen-independent [26, 30]. We also reported that PC3 prostate cancer cells and HEK293 human embryonic kidney cells, both of which lack AR, were less sensitive to growth inhibition upon MED19 depletion compared to LNCaP-abl cells [26].

We expected that the selective effect by overexpressed MED19 on AR-associated gene expression, as well as genome-wide occupancy of MED19, AR, and H327 acetylation under androgen deprivation, would illuminate mechanism. Indeed, we observed by RNA-seq a defined set of genes that were differentially expressed upon MED19 overexpression in LNCaP cells compared to control cells. In addition, we observed a large overlap between MED19 and AR occupancy under both androgen-independent and androgen-dependent conditions. There was also a unique set of loci with AR and MED19 occupancy under androgen deprivation in MED19 LNCaP cells compared to control cells, suggesting that MED19 can drive AR to new sites. Thus, we observed upon MED19 overexpression a selective alteration of the AR cistrome. This was reflected in changes in gene expression under conditions of low androgen levels.

We also found that MAOA was upregulated upon MED19 overexpression in LNCaP cells under androgen deprivation. This was associated with an increase in occupancy of AR and H3K27 acetylation at the MAOA promoter. We also observed a striking growth-inhibitory effect upon MAOA depletion in MED19 LNCaP cells under androgen deprivation. Although MAOA is likely not the sole mediator of MED19-induced androgen-independent growth, it is important, given the large reduction in androgen-independent growth upon its depletion. This is in contrast to the small growth inhibitory effect of LRRTM3 depletion. Furthermore, multiple studies have established the importance of increased MAOA expression in facilitating prostate cancer proliferation [34, 35, 45, 46]. Conversely, a polymorphism in the MAOA promoter conferring low expression is associated with lower risk of developing prostate cancer [47].

An ELK1 motif was enriched at sites occupied by AR, as well as MED19, in MED19 LNCaP cells but not in control LNCaP cells. Furthermore, MED19-upregulated genes, including MAOA, were associated with ELK1, suggesting that ELK1 could be cooperating with MED19 and AR to promote MAOA expression and growth under androgen deprivation. Indeed, MED19 overexpression promoted ELK1 occupancy at the MAOA promoter, and ELK1 depletion reduced MAOA expression and androgen-independent growth. ELK1 is an ETS transcription factor that controls AR transcriptional activity and promotes prostate cancer progression [20, 23, 24]. ELK1 has also been shown to interact directly with the AR ligand-independent N-terminal transcriptional activation domain [23, 24]. Given the increase in H3K27 acetylation at the MAOA promoter with MED19 overexpression, it is possible that MED19, in conjunction with ELK1, could promote recruitment of HATs, such CBP and p300, which are also known AR co-regulators [10–12]. Consistent with this change in H3K27 acetylation, ELK1 has also been found to interact with CBP and p300 in other cell types [48, 49].

Based on our findings, we propose a model whereby under conditions of androgen deprivation and MED19 upregulation, MED19-containing Mediator cooperates with ELK1 to recruit and stabilize AR, via its N-terminal domain, to the promoter of MAOA, and increases H3K27 acetylation at the MAOA promoter, through recruitment of HATs. Recruitment of Pol II ensues, upregulating MAOA and licensing cell growth under low androgen (Fig 9). Consistent with this model, the structural determination of the yeast Mediator complex by cryo-EM revealed MED19 contacts the carboxy terminal domain (CTD) tail of Pol II; these contacts between Mediator and the Pol II CTD serve to recruit and stabilize Pol II [50]. Recent structure determination of the mammalian Mediator complex confirmed the importance of the middle module for CTD contact [27].

**Fig 9.**
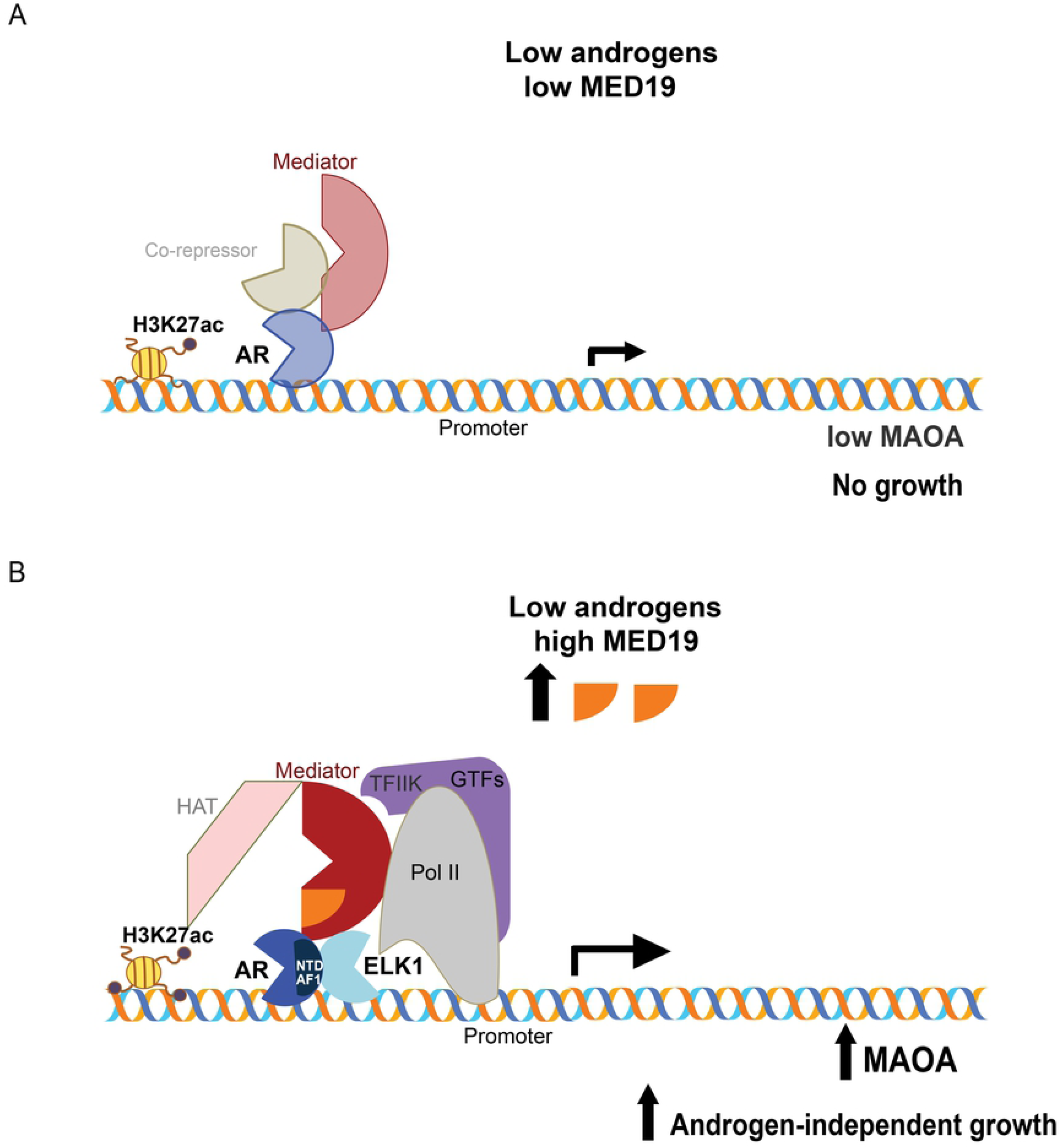
Model of MED19 driving androgen-independent growth by cooperating with ELK1 to promote AR occupancy and H3K27 acetylation at the MAOA promoter. A) Under low androgen and low MED19, AR occupancy is low at the MAOA promoter, MAOA is weakly expressed, and cells are growth-inhibited. B) When MED19 is upregulated, MED19 in Mediator cooperates with ELK1 to recruit and stabilize AR via its N-terminal domain (NTD) at the MAOA promoter, also recruiting Pol II and HATs, upregulating MAOA expression and driving androgen-independent growth.

Another Mediator subunit, MED1 (also a middle subunit), has been described to promote androgen-dependent AR activity through interaction with the ligand-binding domain of AR, and is overexpressed in prostate cancer [51–54]. It is possible that MED1 (or other Mediator subunits) may play a role in MED19-induced androgen-independent growth. Indeed, when each of the 33 subunits of Mediator is depleted under androgen deprivation in both MED19 LNCaP cells and LNCaP-abl cells, depletion of MED1, as well as MED14, MED15, and MED16 (previously reported to affect prostate cancer cell growth and AR transcriptional activity), significantly reduced androgen-independent growth, with an overall variable effect on growth from subunit-to-subunit (S15A and S15B Fig) [26]. Strikingly, no subunit had a significantly greater effect on androgen-independent growth than MED19, highlighting the crucial and specific function of MED19.

Our study may also have implications for MED19 function in other cancers. For example, reduction of MED19 by siRNA has been reported to reduce the proliferation of certain breast, ovarian, cervical, and lung cancer cell lines, and increased abundance of MED19 protein is observed in tumors relative to benign tissue [55–62]. This suggests that MED19 functions with other transcription factors in other cell types to regulate gene expression and enhance cellular proliferation. Although the regulation of gene expression and cancer cell characteristics by Mediator subunits is complex, our study provides key insight into the mechanism of MED19 action in prostate cancer cells and androgen independence.

## Materials and methods

### Generation of LNCaP cells with stable overexpression of MED19

LNCaP cell lines purchased from the ATCC (Manassas, VA) were used for stable transfections. MED19 overexpression and empty vectors were purchased from Origene Technologies (NM_153450; plenti-myc-DDK backbone). Cells were generated after lentiviral infection with the above constructs. Lentiviral particles were produced in the 293T/17 cell line (ATCC). LNCaP, RWPE-1, RWPE-2, and mouse prostate stem cell lines were infected on two consecutive days with control or MED19 lentiviral particles and polybrene. Pooled clones were collected after selection with puromycin (1 μg/mL). MED19 expression was verified by western blot.

### Cell culture and reagents

LNCaP cell lines were maintained in complete RPMI: RPMI-1640 supplemented with 10% fetal bovine serum (FBS) (Hyclone; Fisher Scientific) and 1% penicillin-streptomycin (Cellgro; Mediatech, Inc.) For assays under androgen deprivation, cells were cultured in androgen-depleted RPMI: phenol red- and L-glutamine-free RPMI-1640 supplemented with 10% FBS dextran charcoal-stripped of androgens (c-FBS, Hyclone) and 1% L-glutamine (Cellgro Mediatech, Inc.). Cells were cultured on poly-d-lysine-coated plates. RWPE-1 cells and RWPE-2 cells were cultured in keratinocyte SFM media supplemented with L-glutamine, BPE, and EGF (ThermoFisher Scientific) [63]. The mouse prostate stem cell line expressing activated AKT was cultured as described in [31]. R1881 (Perkin Elmer) was reconstituted in ethanol. Enzalutamide (MedKoo) was reconstituted in DMSO. Puromycin (Sigma Aldrich) was reconstituted in water.

### Proliferation assay

Cells were plated in the appropriate media in quintuplicate (LNCaP: 3,000 cells per well in complete media and 5,000 cells per well in androgen-depleted media; RWPE-1: 2,000 cells per well; RWPE-2: 10,000 cells per well; mouse prostate stem cells: 1,000 cells per well) in poly-d-lysine-coated 96-well plates. Cell proliferation was determined using the Cyquant-NF Cell Proliferation Assay (Invitrogen) or PrestoBlue Cell Viability Assay (ThermoFisher Scientific). Fluorescence was quantified with the SpectraMaxM5 Microplate Reader and SoftMaxPro software (Molecular Devices) and normalized to readings at Day 0 (day after plating).

### Colony formation assay

Cells were plated in the appropriate media in duplicate (LNCaP: 5,000 cells per well in complete media and 10,000 cells per well in androgen-depleted media; RWPE-1, RWPE-2, mouse prostate stem cells: 10,000 cells per well) in poly-d-lysine-coated 6-well plates for 10-14 days. Cells were fixed with 66% methanol/33% acetic acid solution and stained with 0.1% crystal violet solution.

### Spheroid formation assay

Cells were plated in the appropriate media in 96-well ultra-low attachment plates (Corning) (LNCaP: 1,000-2,000 cells per well, 8 wells/condition) for 10-14 days. Cells were imaged with CellInsight CX7 LZR and spheroid area per well was analyzed by Cellomics Scan Version 6.6.1.

### Xenograft study

For xenograft experiments, mouse prostate stem cells (5 × 10^6^) were mixed with an equal volume of Matrigel and injected subcutaneously into the flank region of Nu/J (nude) male mice (Jackson Laboratories). Tumor volume was measured twice weekly. All animal studies were performed at NYU School of Medicine. The animal research was approved by the NYU School of Medicine Institutional Animal Care and Use Committee (IACUC), protocol number IA16-01775.

### RNA preparation and quantitative RT-PCR

Total RNA was extracted using RNeasy (Qiagen) according to the manufacturer’s instructions. RNA (1 μg) was reverse transcribed using the Verso cDNA Synthesis Kit (ThermoFisher Scientific) following the manufacturer’s instructions. Gene-specific cDNA was amplified in a 10-μL reaction containing Fast SYBR Green qPCR Master Mix (ThermoFisher Scientific). Real-time PCR was performed using the Applied Biosystems Quantstudio 6 Flex Real-Time PCR System with each gene tested in triplicate. Data were analyzed by the DDCT method using RPL19 as a control gene, and normalized to control samples, which were arbitrarily set to 1. The sequences of the primers used for real-time PCR are as follows. RPL19 – F:CACAAGCTGAAGGCAGACAA, R:GCGTGCTTCCTTGGTCTTAG; ELK1 – F:CACATCATCTCCTGGACTTCAC, R:CGGCTGAGCTTGTCGTAAT; MED19 – F:CTGTGGCCCTTTTTACCTCA, R:GCTTCTCCTTCACCTTCTTCC; AR – F:TACCAGCTCACCAAGCTCCT, R:GAACTGATGCAGCTCTCTCG; LRRTM3 – F:ATACGACCAGCCCACAATAAG, R: GCTCAGTCTCTAGGTGTGTTTC; MAST4 – F:GCCAAAGAAGGACAGGGTATTA, R:GCTGTCCCACTATCGTAGTTTC; MAOA – F:CCTGTGGTTCTTGTGGTATGT, R:CACCTACAAACTTCCGTTCCT; AR-V7 – F: CCATCTTGTCGTCTTCGGAAATGTTA, R:TTTGAATGAGGCAAGTCAGCCTTTCT; FKBP5 – F:CGCAGGATATACGCCAACAT, R:CTTGCCCATTGCTTTATTGG; PSA (KLK3) – F:CCAAGTTCATGCTGTGTGCT, R:GCACACCATTACAGACAAGTGG.

### SiRNA knockdowns

Three individual siRNAs (Silencer Select; Ambion, Life Technologies) were pooled and transfected into cells using the Lipofectamine RNAimax transfection reagent (ThermoFisher Scientific) following the manufacturer’s instructions. Nonsilencing (scrambled) siRNAs were used as controls. siRNAs were used at a final concentration of 25 nM.

### Protein extraction and Western blot analysis

Cells were lysed in RIPA buffer supplemented with protease inhibitor cocktail (Cell Signaling Technology). Protein lysates were subjected to SDS/PAGE and immunoblotted with antibodies against AR (441, Santa Cruz Biotechnology; cat # sc-7305) or MYC tag (Cell Signaling Technology; cat **#** 2276S). Tubulin (Covance; cat # MMS-489P) was used as a loading control.

### RNA-sequencing

RNA was prepared as described above. Libraries were prepared with ribodepletion using Illumina TruSeq stranded total RNA with RiboZero Gold library preparation kit. Sequencing was performed using the Illumina HiSeq2500 Sequencing system (HiSeq 4000 Paired-End 50 or PE75 Cycle Lane). Data was analyzed by Rosalind (https://rosalind.onramp.bio/), with a HyperScale architecture developed by OnRamp BioInformatics, Inc. (San Diego, CA). Reads were trimmed using cutadapt. Quality scores were assessed using FastQC. Reads were aligned to the Homo sapiens genome build hg19 using STAR. Individual sample reads were quantified using HTseq and normalized via Relative Log Expression (RLE) using DESeq2 R library. Read Distribution percentages, violin plots, identity heatmaps, and sample MDS plots were generated as part of the QC step using RSeQC. DEseq2 was also used to calculate fold changes and p-values. Clustering of genes for the final heatmap of differentially expressed genes was done using the PAM (Partitioning Around Medods) method using the fpc R library. Functional enrichment analysis of pathways, gene ontology, domain structure and other ontologies was performed using HOMER. Several database sources were referenced for enrichment analysis, including Interpro, NCBI, MSigDB REACTOME, WikiPathways. Enrichment was calculated relative to a set of background genes relevant for the experiment. Additional gene enrichment is available from the following partner institutions: Advaita (http://www.advaitabio.com/ipathwayguide). Differentially expressed genes were also analyzed using the Enrichr platform from the Ma’ayan Laboratory, including ChEA and “Transcription Factor (TF) Perturbations Followed by Expression” analyses [64, 65].

### Chromatin immunoprecipitation (ChIP)-sequencing

Cells were double crosslinked with formaldehyde and the bifunctional protein crosslinker disuccinimidyl glutarate (DSG) to preserve both protein-DNA and protein-protein interactions. A mixture of two antibodies for AR ChIP was used, one against the C-terminus and one against the N-terminus, to maximize AR enrichment and minimize epitope masking. Antibodies were tested and optimized. The ChIP-seq study was performed in independent biological triplicates (one sample for ChIP-seq for AR in control LNCaP cells + R1881 was excluded from the analyses because of low signal.) Inputs were used for normalization and additional IgG controls included to ensure any low occupancy peaks were specific to AR and not background. We discarded any peaks for AR and for H3K27 acetylation that scored above 0 for IgG; this was also done with peaks for FLAG-MED19 in MED19 LNCaP cells that scored above 0 for FLAG in control LNCaP cells (very few peaks scored above 0 and overlapped in either case) (S6C Fig and S7 Table). Rigorous scoring of peaks was done using the Rosalind platform (see below).

Protocol for ChIP-seq was adapted from Fonseca *et al* [66]. Briefly, cells were double cross-linked with DSG (ProteoChem; cat # c1104) for 20 min and 1% formaldehyde for 10 min. Crosslinking was quenched with Tris-HCl pH 7.5 (Invitrogen). Cells were collected, washed with PBS, and cell pellets snap frozen with liquid nitrogen. Cell pellets were resuspended in nuclei isolation buffer (50 mM Tris–pH 8.0, 60 mM KCl, 0.5% NP40), nuclei collected, and resuspended in sonication buffer (RIPA buffer). Samples were sonicated in TPX PMP tubes (Diagenode) for 60 min (30 sec. on, 30 sec. off) in a Bioruptor sonicator (Diagenode). Inputs (10%) were collected and supernatants were then incubated overnight with the following antibodies pre-incubated with Protein A and Protein G Dynabeads (Invitrogen): a mixture of AR C-terminal (441, Santa Cruz Biotechnology, cat # sc-7305) and AR N-terminal (Cell Signaling Technology, cat # 5153); DYKDDDDK Tag (FLAG epitope) (Cell Signaling Technology, cat # 14793); or H3K27 acetylation (Active Motif, cat # 39034). Control ChIPs were performed with normal mouse IgG (Santa Cruz Biotechnology, cat # sc-2025) and normal rabbit IgG (Sigma Aldrich, cat # 12-370). Immunocomplexes were then washed and cross-linking reversed overnight at 65 °C with 5 M NaCl. DNA was isolated with the Zymo Chip DNA Clean and Concentrator kit (Zymo Research). Libraries were prepared according to the protocol described in [66]. Sequencing was performed using Illumina HiSeq4000 Sequencing (HiSeq 4000 Single Read 50 Cycle Lane).

Data were analyzed by Rosalind (https://rosalind.onramp.bio/), with a HyperScale architecture developed by OnRamp BioInformatics, Inc. (San Diego, CA). Reads were trimmed using cutadapt. Quality scores were assessed using FastQC. Reads were aligned to the Homo sapiens genome build hg19 using bowtie2. Per-sample quality assessment plots were generated with HOMER and Mosaics. Peaks were called using MACS2 (with input controls background subtracted). Peak overlaps were analyzed using the DiffBind R library. Read distribution percentages, identity heatmaps, and FRiP plots were generated as part of the QC step using ChIPQC R library and HOMER. HOMER was also used to generate known and de novo motifs and perform functional enrichment analysis of pathways, gene ontology, domain structure and other ontologies.

### ChIP-qPCR

ChIPs were performed as described above, with ChIPs for AR, FLAG epitope, H3K27 acetylation, and ELK1 (abcam, cat # ab32106). IgGs were included as negative controls. After DNA isolation, qPCR was performed as described above, with primers targeting the MAOA promoter region - F: TGTCAAGGCAGGCGTCTAC, R: GGACCCTTGTACTGACAC. Relative enrichment was calculated as a percentage of 10% input.

### Statistical analyses

Statistical analyses were performed using GraphPad Prism software. Data are reported as mean ± SEM (technical replicates for each experiment described above). Number of experiments are described in the figure legends; unless otherwise noted, two-tailed unpaired Student’s t test was used when comparing two groups, with a p value < 0.05 being considered significant and levels of significance denoted as *p < 0.05; **p < 0.01; and ***p < 0.001.

## Acknowledgements

We thank Dr. Chi Yun, Dr. Sokha Nhek, Rebecca Lee, Dr. David Kahler, and the NYU High Throughput Biology Laboratory for technical support and advice. We thank Dr. Adriana Heguy, Paul Zappile, and the Genome Technology Center for technical support. We thank Drs. Christopher Glass, Gregory Fonseca, and Jenhan Tao for reagents, protocols, and technical assistance with the sequencing studies. We thank Dr. Elaine Wilson for the AKT-transformed mouse prostate stem cell line. We thank Dr. Gregory David and Susan Ha for critical insight and assessment of the manuscript, and the Garabedian and Logan laboratories for their support.

## Author contributions

**Project administration:** Michael J. Garabedian

**Funding acquisition:** Michael J. Garabedian

**Supervision:** Michael J. Garabedian

**Conceptualization:** Michael J. Garabedian, Hannah Weber

**Investigation:** Hannah Weber, Rachel Ruoff

**Formal analysis:** Hannah Weber, Rachel Ruoff, Michael J. Garabedian

**Validation:** Hannah Weber, Rachel Ruoff

**Visualization:** Hannah Weber, Rachel Ruoff

**Writing – original draft preparation:** Hannah Weber

**Writing – review & editing:** Rachel Ruoff, Michael J. Garabedian

## Supporting information

**S1 Fig. MED19 LNCaP cells stably overexpress MED19**. MED19 LNCaP cells with stable overexpression of MYC- and FLAG-tagged MED19 and control LNCaP cells with stable expression of the empty vector were created by lentiviral transduction, with pooled clones selected with puromycin. After selection, stable overexpresson of MED19 in MED19 LNCaP cells was confirmed. A) RNA was extracted and qPCR was performed for MED19 in control LNCaP cells and MED19 LNCaP cells to confirm upregulation of MED19 mRNA (fold change expression normalized to RPL19 with MED19 mRNA expression in control LNCaP cells set as “1”). *p < 0.05; **p < 0.01; and ***p < 0.001. B) MED19 LNCaP cells and control LNCaP cells were treated with MED19 siRNA or scrambled siRNA, and total protein lysates were probed by MYC tag. Tubulin was used as a loading control.

**S2 Fig**. **MED19 overexpression in RWPE-1 cells and mouse prostate stem cells promotes colony formation**. A) RWPE-1, B) RWPE-2, or C) mouse stem cells with activated AKT (MSC), stably overexpressing MED19 (MED19 RWPE-1/RWPE-2/MSC) or control empty vector (control RWPE-1/RWPE-2/MSC), were cultured in their standard media. Colony formation was evaluated by culturing the cells at low density for 14 days and fixing and staining with crystal violet. Experiments were performed in biological duplicate, with representative results shown.

**S3 Fig. MED19 RWPE-1 cells have comparable MED19 expression to MED19 RWPE-2 cells**. Total protein lysates from RWPE-1 and RWPE-2 cells stably expressing FLAG- and MYC-tagged MED19 (MED19 RWPE-1 and MED19 RWPE-2) or empty vector (control RWPE-1 and control RWPE-2) were probed for MYC tag, with tubulin used a loading control.

**S4 Fig. MED19 LNCaP cells do not express AR-V7**. MED19 LNCaP cells and control LNCaP cells were cultured under androgen deprivation for 3 days and treated overnight with ethanol vehicle. RNA was extracted and mRNA measured by qPCR for AR-V7 mRNA (fold change expression normalized to RPL19 with AR-V7 mRNA expression in control LNCaP cells set as “1”). LNCaP-95 cells that express AR-V7 were used as a positive control. *p < 0.05; **p < 0.01; and ***p < 0.001.

**S5 Fig. MED19 LNCaP cells are sensitive to AR knockdown**. MED19 LNCaP cells were cultured in A) androgen-depleted media or B) androgen-containing media, with control LNCaP cells. AR was depleted by siRNA and proliferation was evaluated after 7 days, normalized to proliferation with scrambled siRNA. KIF11 was used as a positive control. Experiment was performed in biological duplicate, with representative results shown. *p < 0.05; **p < 0.01; and ***p < 0.001.

**S6 Fig. QC of ChIP-seq samples**. MED19 LNCaP cells and control LNCaP cells were cultured under androgen deprivation for 3 days and treated overnight (16 h) with ethanol vehicle or 10 nM R1881. ChIP-seq for FLAG-MED19, AR, and H3K27ac was performed in biological triplicate. A) ChIP-qPCR QC of AR, H3K27ac, and FLAG-MED19 ChIPs are shown, with normalization to inputs. AR occupancy and H3K27ac at PSA ARE III greatly increase in response to R1881 treatment. IgG is shown as a negative control. FLAG-MED19 shows high occupancy in MED19 LNCaP cells at PDZK1P1, identified as a site of strong FLAG-MED19 occupancy from a pilot ChIP-seq for FLAG-MED19. FLAG in control LNCaP cells is shown as a negative control. Experiments were performed in biological triplicate, with representative results shown. *p < 0.05; **p < 0.01; and ***p < 0.001. B) ChIP-seq tracks (representative result) for AR and H3K27ac at PSA and FKBP5 in control LNCaP cells with vehicle or R1881 treatment. AR occupancy and H3K27ac clearly increase in response to R1881 treatment (occupancy scores in S7 Table). C) Overlap of IgG with AR (left) and H3K27ac sites (middle); and overlap of FLAG in control LNCaP cells with FLAG-MED19 in MED19 LNCaP cells (right). All normalized to input. IgG and FLAG-control yield very few sites, with minimal overlap (all sites in S7 Table).

**S7 Fig. QC and qPCR validation of RNA-seq**. MED19 LNCaP cells and control LNCaP cells were cultured under androgen deprivation for 3 days and treated overnight (16 h) with ethanol vehicle or 10 nM R1881. RNA-seq was performed in biological triplicate. Graphs represent fold changes from qPCR (fold change expression normalized to RPL19 with PSA or FKBP5 mRNA expression in vehicle-treated cells set as “1”) and tables represent fold changes from RNA-seq. Upregulation of PSA and FKBP5 mRNA expression in A) control LNCaP cells and B) MED19 LNCaP cells in response to R1881 treatment, with consistency between RNA-seq and qPCR, and expected increase in expression with R1881 treatment. Experiments were performed in biological triplicate, with representative results shown. *p < 0.05; **p < 0.01; and ***p < 0.001.

**S8 Fig. MED19 alters mRNA expression, AR occupancy, and H3K27 acetylation for LRRTM3 +/- androgens, but LRRTM3 does not affect androgen-independent growth**. A, B) MED19 LNCaP cells and control LNCaP cells were cultured under androgen deprivation for 3 days and treated overnight (16 h) with ethanol vehicle or 10 nM R1881. RNA-seq and ChIP-seq for FLAG-MED19, AR, and H3K27ac were performed in biological triplicate, with the exception of ChIP-seq for AR in control LNCaP cells + R1881, where one sample was excluded from the analyses because of low signal. A) Fold change mRNA expression from RNA-seq and qPCR validation of changes of LRRTM3 mRNA expression (performed in biological triplicate, representative result shown; fold change expression normalized to RPL19 with LRRTM3 mRNA expression in vehicle-treated control LNCaP cells set as “1”). Greater fold changes by qPCR likely due to low abundance (raw counts in RNA-seq) of LRRTM3 in control LNCaP cells. B) ChIP-seq tracks (representative results) for FLAG-MED19, AR, and H3K27ac for androgen deprivation or R1881 treatment are shown for intronic regions of LRRTM3. Fold change (up (+) or down (-)) in occupancy scores for MED19 LNCaP cells compared to control LNCaP cells shown for each peak (see S6 Table for all occupancy scores). ++ indicates positive occupancy score in MED19 LNCaP cells and a score of zero in control LNCaP cells; -- indicates an occupancy score of zero in MED19 LNCaP cells and a positive score in control LNCaP cells. C) LRRTM3 was depleted by siRNA and proliferation of MED19 LNCaP cells in androgen-depleted media was evaluated after 7 days, normalized to proliferation with scrambled siRNA (negative control, light grey). KIF11 knockdown is included as a positive control (black). Experiment was performed in biological duplicate, with representative results shown. *p < 0.05; **p < 0.01; and ***p < 0.001.

**S9 Fig. MED19 alters mRNA expression, AR occupancy, and H3K27 acetylation for MAST4**. A,B) MED19 LNCaP cells and control LNCaP cells were cultured under androgen deprivation for 3 days and treated overnight (16 h) with ethanol vehicle or 10 nM R1881. RNA-seq and ChIP-seq for FLAG-MED19, AR, and H3K27ac were performed and in biological triplicate., with the exception of ChIP-seq for AR in control LNCaP cells + R1881, where one sample was excluded from the analyses because of low signal. A) Fold change mRNA expression from RNA-seq and qPCR validation of changes of MAST4 mRNA expression (performed in biological triplicate, representative result shown; fold change expression normalized to RPL19 with MAST4 mRNA expression in vehicle-treated control LNCaP cells set as “1”). *p < 0.05; **p < 0.01; and ***p < 0.001. B) ChIP-seq tracks (representative results) for FLAG-MED19, AR, and H3K27ac for androgen deprivation or R1881 treatment are shown for promoter and intronic regions of MAST4. Fold change (up (+) or down (-)) in occupancy scores for MED19 LNCaP cells compared to control LNCaP cells shown for each peak (see S6 Table for all occupancy scores). ++ indicates positive occupancy score in MED19 LNCaP cells and a score of zero in control LNCaP cells; -- indicates an occupancy score of zero in MED19 LNCaP cells and a positive score in control LNCaP cells.

**S10 Fig. MED19 and androgen treatment promote mRNA expression, AR occupancy, and H3K27 acetylation for MAOA**. A, B) MED19 LNCaP cells and control LNCaP cells were cultured under androgen deprivation for 3 days and treated overnight (16 h) with ethanol vehicle or 10 nM R1881. RNA-seq and ChIP-seq for FLAG-MED19, AR, and H3K27ac were performed in biological triplicate, with the exception of ChIP-seq for AR in control LNCaP cells + R1881, where one sample was excluded from the analyses because of low signal. A) Fold change mRNA expression from RNA-seq and qPCR validation of changes of MAOA mRNA expression (performed in biological triplicate, representative result shown; fold change expression normalized to RPL19 with MAOA mRNA expression in vehicle-treated control LNCaP cells set as “1”). B) ChIP-seq tracks (representative results) for FLAG-MED19, AR, and H3K27ac for androgen deprivation or R1881 treatment are shown for promoter region of MAOA. Fold change (up (+) or down (-)) in occupancy scores for MED19 LNCaP cells compared to control LNCaP cells shown for each peak (see S6 Table for all occupancy scores). ++ indicates positive occupancy score in MED19 LNCaP cells and a score of zero in control LNCaP cells; -- indicates an occupancy score of zero in MED19 LNCaP cells and a positive score in control LNCaP cells. C) ChIP-qPCR for FLAG-MED19, AR, and H3K27ac at the MAOA promoter overlapping with published ARE. (Control - control LNCaP cells; MED19 – MED19 LNCaP cells; veh – vehicle treatment; R1881 – R1881 treatment). Experiment was performed in duplicate, with representative results shown. *p < 0.05; **p < 0.01; and ***p < 0.001.

**S11 Fig. FOXA1 and FOXM1 are enriched at sites of AR occupancy and MED19 occupancy in the absence and presence of androgens**. MED19 LNCaP cells and control LNCaP cells were cultured under androgen deprivation for 3 days and treated overnight (16 h) with ethanol vehicle or 10 nM R1881. ChIP-seq for FLAG-MED19, AR, and H3K27ac was performed in biological triplicate. A) Top 10 enriched transcription factor motifs under androgen deprivation, associated with MED19 sites in MED19 LNCaP cells (top), MED19 and AR occupied sites in MED19 LNCaP cells (middle top), AR sites in MED19 LNCaP cells (middle bottom), and AR sites in control LNCaP cells (bottom). B) Top 10 enriched transcription factor motifs with R1881 treatment, associated with MED19 sites in MED19 LNCaP cells (top), MED19 and AR occupied sites in MED19 LNCaP cells (middle top), AR sites in MED19 LNCaP cells (middle bottom), and AR sites in control LNCaP cells (bottom).

**S12 Fig. Enriched ChIP-seq motifs unique to AR+MED19 in MED19 LNCaP cells with R1881 treatment and gene changes associated with Enrichr Transcription Factor Perturbation with vehicle or R1881 treatment.** A) Top 10 enriched transcription factor motifs with R1881 treatment associated with sites of AR and MED19 occupancy in MED19 LNCaP cells where AR is present only in MED19 LNCaP cells, with SP1 as the top associated transcription factor. B) Gene changes associated with ELK1 knockdown and AR knockdown from Enrichr Transcription Factor Perturbation, compared to MED19 overexpression under androgen deprivation from the RNA-seq study. C) SRF knockdown is the top hit from Enrichr Transcription Factor Perturbation, associated with genes upregulated by MED19 overexpression with R1881 treatment from the RNA-seq study (top); corresponding genes changes associated with SRF knockdown compared to MED19 overexpression (bottom).

**S13 Fig**. **FOXA1 and FOXM1 and AR-related motifs are enriched at sites of AR and MED19 occupancy with R1881 treatment**. MED19 LNCaP cells and control LNCaP cells were cultured under androgen deprivation for 3 days and treated overnight (16 h) with ethanol vehicle or 10 nM R1881. ChIP-seq for FLAG-MED19, AR, and H3K27ac was performed in biological triplicate. A) Top 10 enriched transcription factor motifs associated with AR sites in control LNCaP cells in R1881 vs. vehicle treatment, with enrichment of AR-related motifs in response to R1881 treatment. B) Top 10 enriched transcription factor motifs associated with AR sites in MED19 LNCaP cells in R1881 vs. vehicle treatment, with enrichment of AR-related motifs in response to R1881 treatment. C) Top 10 enriched transcription factor motifs associated with MED19 sites in MED19 LNCaP cells in R1881 vs. vehicle treatment, with enrichment of AR-related motifs in response to R1881 treatment.

**S14 Fig. QC of ELK1 ChIP**. MED19 LNCaP cells and control LNCaP cells were cultured under androgen deprivation for 3 days and treated overnight (16 h) with ethanol vehicle or 10 nM R1881, and ChIP-qPCR for ELK1 was performed. ELK1 occupancy at previously published ELK1 sites was verified in control LNCaP cells (top) and MED19 LNCaP cells (bottom), with occupancy verified +/- R1881 for sites at A) Chr.1 and B) Chr. 6. Normalization to inputs was done and IgG is shown as a negative control. *p < 0.05; **p < 0.01; and ***p < 0.001.

**S15 Fig. Depletion of Mediator subunits in MED19 LNCaP cells and LNCaP-abl cells under androgen deprivation.** Each Mediator subunit or associated factor from the kinase module was depleted by siRNA and proliferation in androgen-depleted media was evaluated after 5 days, normalized to proliferation with scrambled siRNA (negative control, light grey). KIF11 knockdown is included as a positive control (black). MED19 depletion is highlighted in bold. A) Knockdown of Mediator subunits in MED19 LNCaP cells. B) Knockdown of Mediator subunits in LNCaP-abl cells. *p < 0.05; **p < 0.01; and ***p < 0.001. Statistics denote comparison to scrambled siRNA. There is no statistically significant difference in growth between MED19 depletion and MED1 depletion in MED19 LNCaP cells or in LNCaP-abl cells. There is no statistically significant difference in growth between MED19 depletion and MED26/MED4/MED18/CDK19/MED12/MED27 depletion in MED19 LNCaP cells.

**S16 Fig. Full western blot for MED19 LNCaP cell stable overexpression of MED19 protein from S1 Fig**. A) Full western blot from Supplemental Figure S1. A) Membrane overlay of full western blot from Supplemental Figure S1.

**S1 Table. Full list of 151 genes from RNA-seq significantly altered ≥1.5-fold (p-adj≤0.05) in MED19 LNCaP cells compared to control LNCaP cells, cultured under androgen deprivation for 3 days and treated overnight (16 h) with ethanol vehicle.** 76 genes are upregulated (including MED19, top) and 75 genes are downregulated. P-values, p-adjusted values, fold changes, and gene descriptions are shown for each gene (sheet 1). Genes responsive to R1881 treatment from the RNA-seq (sheet 2), AR targets from ChEA (sheet 3), and androgen-responsive or AR targets from the literature (sheet 4) are shown.

**S2 Table. Full list of 309 genes from RNA-seq significantly altered ≥1.5-fold (p-adj≤0.05) in MED19-LNCaP cells compared to control LNCaP cells, cultured under androgen deprivation for 3 days and treated overnight (16 h) with 10 nM R1881.** 78 genes are upregulated (including MED19, top) and 231 genes are downregulated. P-values, p-adjusted values, fold changes, and gene descriptions are shown for each gene (sheet 1). Genes responsive to R1881 treatment from the RNA-seq (sheet 2), AR targets from ChEA (sheet 3), and androgen-responsive or AR targets from the literature (sheet 4), are shown.

**S3 Table. Full list of 4430 genes from RNA-seq significantly altered ≥1.5-fold (p-adj≤0.05) in control LNCaP cells treated overnight (16 h) with 10 nM R1881 compared to control LNCaP cells treated overnight (16 h) with ethanol vehicle, under androgen deprivation for 3 days.** 2361 genes are upregulated and 2069 genes are downregulated. P-values, p-adjusted values, fold changes, and gene descriptions are shown for each gene.

**S4 Table. Full list of 5041 genes from RNA-seq significantly altered ≥1.5-fold (p-adj≤0.05) in MED19 LNCaP cells treated overnight (16 h) with 10 nM R1881 compared to MED19 LNCaP cells treated overnight (16 h) with ethanol vehicle, under androgen deprivation for 3 days.** 2727 genes are upregulated and 2314 genes are downregulated. P-values, p-adjusted values, fold changes, and gene descriptions are shown for each gene.

**S5 Table. Response of MED19 LNCaP cells vs. control LNCaP cells to androgens**. Top 100 androgen-induced (sheet 1) and androgen-repressed (sheet 2) genes for control LNCaP cells and MED19 LNCaP cells, with fold changes for each shown in comparison. Genes with 1.5-fold or more change in mRNA expression in response to androgen only in control LNCaP cells (sheet 3) or only in MED19 LNCaP cells (sheet 4).

**S6 Table. FLAG-MED19, AR, and H3K27ac occupancy in MED19 LNCaP cells at genes differentially expressed with MED19 overexpression in the absence and presence of androgens.** Occupancy shown for androgen deprivation (sheet 1) and with R1881 treatment (sheet 2), with fold change in expression upon MED19 overexpression indicated. Androgen responsiveness of each gene in control LNCaP cells and MED19 LNCaP cells is also indicated. AR occupancy scores for LRRTM3 (sheet 3), MAST4 (sheet 4), and MAOA (sheet 5).

**S7 Table. AR occupancy score QC and background peaks.** AR occupancy scores for PSA and FKBP5 (sheet 1); and list of background peaks for IgG (sheet 2) and FLAG in control LNCaP cells (sheet 3) for ChIP-seq.

